# Systems Biology Approach to Model the Life Cycle of *Trypanosoma cruzi*

**DOI:** 10.1101/025510

**Authors:** Alejandra Carrea, Luis Diambra

**Author notes:** Corresponding author: (LD).

## Abstract

Due to recent advances in reprogramming cell phenotypes, many efforts have been dedicated to developing reverse engineering procedures for the identification of gene regulatory networks that emulate dynamical properties associated with the cell fates of a given biological system. In this work, we propose a systems biology approach for the reconstruction of the gene regulatory network underlying the dynamics of the *Trypanosoma cruzi*’s life cycle. By means of an optimisation procedure, we embedded the steady state maintenance, and the known phenotypic transitions between these steady states in response to environmental cues, into the dynamics of a gene network model. In the resulting network architecture we identified a small subnetwork, formed by seven interconnected nodes, that controls the parasite’s life cycle. The present approach could be useful for better understanding other single cell organisms with multiple developmental stages.

**Abbreviations:**

GRN: gene regulatory network
SVD: singular value decomposition
TS: *trans*-sialidase

## 1 Introduction

One of the main aims in the post-genome era is to elucidate the complex webs of interacting genes and proteins underlying the establishment and maintenance of cell states. Consequently, many researchers have focused on developing quantitative frameworks to identify modules that govern the transitions between different phenotypes. The gene regulatory network (GRN) approach is one of the most popular frameworks used today. This approach has been used to study key reprogramming genes and cell differentiation processes in stem cells from different points of view [1-5]. Mathematically, GRN models are dynamical systems whose states determine the gene-expression levels. The structure of the network is defined as a graph whose nodes are associated with genes (or groups of genes), and whose edges represent the interactions between the nodes. The task of uncovering the GRN architecture from the cell states (gene-expression profiles) represents a very complex inverse problem that has become central in functional genomics [6]. The main drawbacks of this reverse engineering task are not only the large number of genes and the limited amount of data available, but also the nonlinear dynamics of regulations, the inherent experimental errors, the noisy readouts of expression levels, and many other unobserved factors that are part of the challenge. Although emerging technologies offer new prospects for monitoring mRNA concentrations, researchers have focused on determining the architecture of simplified theoretical models.

In this work, we have implemented a GNR approach to analyse transcriptional data of the steady states of the flagellated protozoan parasite *Trypanosoma cruzi (T. cruzi*). This trypanosomatid is the causative agent of Chagas disease, that affects about 7-8 million people worldwide causing about 12,000 deaths per year. Usually, the parasites are transmitted to humans and to other mammalian hosts mainly by contact with the faeces of infected blood-sucking triatomine bugs. *T. cruzi* has several developmental stages both in insect vectors and in mammalian hosts (Fig. 1a). Insects become infected by sucking blood from mammals with circulating parasites (trypomastigotes). In the midgut of the insect, trypomastigotes differentiate into epimastigotes that replicate by binary fission. Then, epimastigotes differentiate into metacyclic trypomastigotes in the hindgut. This parasite form is released in the insects faeces and enters the mammalian host. In turn, metacyclic trypomastigotes invade local cells and differentiate into amastigotes that replicate by binary fission. They subsequently transform into trypomastigotes inside the cells. By lysing the cells, trypomastigotes are released into the circulation. Thus, they spread via the bloodstream and infect new cells from distant tissues where they transform back into intracellular amastigotes. The cycle of transmission is completed when circulating trypomastigotes are taken up in blood meals by triatomine vectors [7].

**Figure 1:**
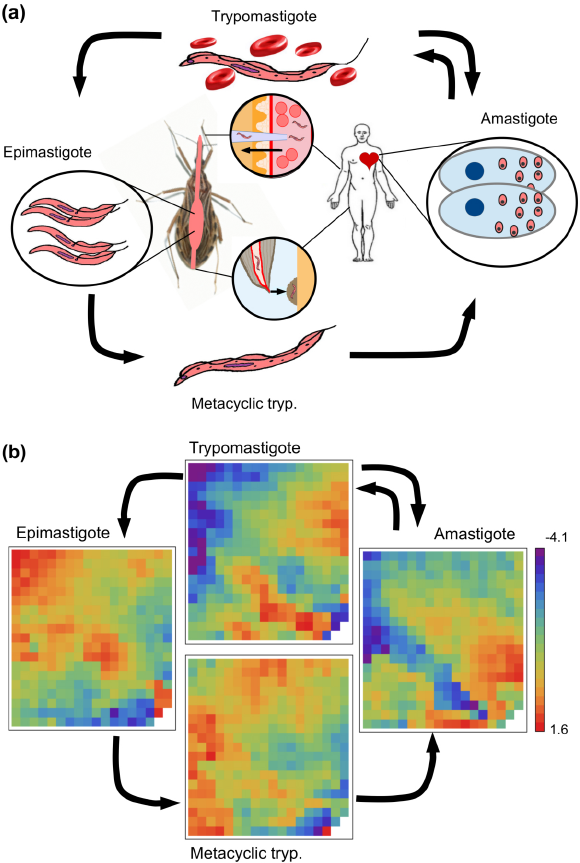
The life cycle of T. *cruzi.* (*a*) Sketch illustrating the life cycle of the parasite. (*b*) Plots illustrating the transcriptional snapshots of the parasite’s four stages. After a dimensional reduction analysis of the microarray dataset, we have found that the four steady states can be represented by 339 variables. Each of these variables (cells in the 19×18 array) corresponds to the intra-cluster average of the log-transformed relative expression level of the genes that belong to the corresponding cluster. Since gene assignment to the clusters is the same for all states, the arrays can be directly compared with one another.

Even though there are potential vaccine candidates against *T.cruzi* infection, no vaccine is yet available [8]. Thus, the finding of novel therapeutic targets remains a significant challenge in the control of Chagas disease. Taking this into consideration, we have implemented a GRN approach to analyse transcriptional data of *T. cruzi*’s steady states [9]. In this framework, we have uncovered the underlying architecture network that supports the steady states associated with the four phenotypic stages of *T. cruzi* and the transitions between the parasite’s life cycle stages in response to environmental cues. We believe that this gene network model can clarify the signaling pathways, predict the response of cellular systems to multiple perturbations other than the ones used to derive the model, and determine the perturbation pattern for any desired response.

## 2 Methods

### 2.1 Microarray data normalisation

In this work we have used the microarray experiments of Minning *et al.* [9]. These data are publicly available in Gene Expression Omnibus (GEO) database (Accession no.: GSE14641). This series is the result of dye-swap experiments, out of which we selected the probe intensity signals of 12 microarrays (three biological replicates, and the four not-mixed parasite stages). These microarrays comprise 12,288 unique 70-mers designed against open reading frames in the annotated CL Brener reference genome sequence. They also contain 500 control oligonucleotides designed from *Arabidopsis* sequences. All of these oligonucleotides were printed in duplicate. Further details about probe preparation, microarray hybridisation, and data acquisition can be found in [9], while the description of the microarrays is available at http://pfgrc.tigr.org.

The probe intensity signals from the microarrays were subjected to the following normalisation procedure. (i) The signal intensity of each probe was set at the average of the signal intensities associated with a pair of replicate spots. (ii) The signal intensity of a probe i was normalised against the average signal of control *Arabidopsis* probes in order to obtain a signal relative intensity within the slide. The average signal of control *Arabidopsis* probes is the arithmetic mean of a set of control probes with valid signals. Of course, this set is the same in all microarray experiments. This normalisation procedure has allowed us to integrate the expression data of all the microarrays. The signal relative intensity of the probe i recorded in one of the biological replicates, *j* = 1, 2, 3, at one of the stages, **α** = 1, 2, 3, 4, was represented by 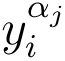. (iii) After within-slide replicates processing, we averaged the relative intensity 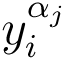 over all replicates belonging to the same stage, i.e., 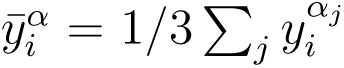 Probes without a valid relative signal in all three biological replicates were not considered in the subsequent analyses. As a result of this processing, we obtained the relative intensity of 8904 probes at parasite’s four different stages. We then considered the variable that describes the expression level of the probe *i* at stage α as the quantity 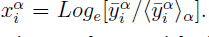 S1 Table lists all normalised expression levels, 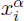, used in the following analyses, with their corresponding oligo IDs.

### 2.2 Clustering procedure

Instead of using each gene’s profile, many researchers have analysed the cell at a higher level of abstraction. One way to do this is by grouping redundant genes, i.e. by clustering co-expressed genes [10,11], and using the average within each cluster as a variable. In order to group the genes by similar activity profiles, we have applied an agglomerative hierarchical clustering method; the Unweighted Pair Group Method with Arithmetic Mean (UPGMA). The agglomerative process is stopped at a given number of clusters considered suitable for our dataset. Since the suitable number of clusters, *N*_*c*_, is not known, it has to be computed beforehand. In order to do this, we repeated the clustering procedure for several *N*_*c*_ values, and computed the Davies-Bouldin index (DBI) as a measure of the clustering merit [11,12]. The DBI is defined as:

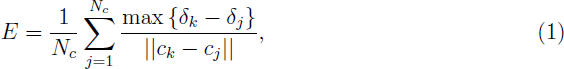

where 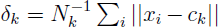 denotes the centroid intra-cluster distances of cluster *k* (*N*_*k*_ being the number of genes belonging to cluster *k*); and 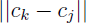 is the distance between the cluster centroids. A low DBI value indicates a good cluster structure. It should be noted that increasing *N*_*c*_ without penalty will always reduce the resulting index. Then, the choice of *N*_*c*_ will intuitively strike a balance between the data compression and the accuracy of the dimensionality reduction. S1 Fig. displays the DBI as a function of *Nc* for the gene-expression profile under study. It can be seen that the DBI does not suffer a significant reduction beyond *N*_*c*_ = 339. Thus, *N*_*c*_ = 339 was selected as the optimal number of clusters. The profiles of the 8,904 genes were grouped in 339 clusters, and the intra-cluster average of the expression level (i.e. 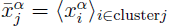) was used in all of the subsequent analyses. Fig. 1b displays a 2D array of the resulting average levels after the dimension reduction process described above for the four stages of *T. cruzi*’s life cycle. The sets of genes belonging to each cluster are listed in S2 Table, and the intra-cluster averages of the expression levels, 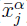, for each cluster (rows) at each of the parasite’s stages (columns) are listed in S3 Table.

### 2.3 Reverse engineering methods

**Gene network dynamics**. In this work, we have implemented a discrete-time linear model [13-15] which has two advantages: it can take into account fluctuations, and its parameter estimation does not involve intensive computational steps [16]. In this model, the system’s state at time *t* is represented by an *N*-dimensional vector *x* (*t*), which represents the activity of the *N* nodes of the network. The temporal evolution of the gene network is governed by:

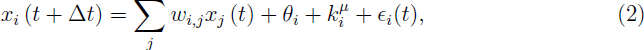

where *w*_*i,j*_ are the elements of the weighted connectivity matrix **W**, *θ*_*i*_ is a constant bias term of gene *i*, and 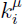 determines the influence of the environmental cue *μ* on gene *i*. We have considered four different cues corresponding to unknown external differentiation signals. Thus, *μ* = 1, 2, 3, and 4 represent the external signals responsible for the transitions to the amastigote, epimastigote, metacyclic tryp., and trypomastigote stages, respectively. *∊_i_*(*t*) is a noise term assumed to be Gaussian with mean equal to 0.

In order to simplify the notation for the parameter estimation procedure, we noticed that the bias term and the environmental cues can be included in an extended version of matrix **W** and of state vector x. Thus, the state of gene *i* is given by:

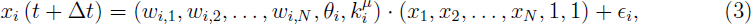

where *μ* corresponds to the acting environmental cue. This said, the same parameter estimation method can be applied whether the environmental cues are present or not.

**Singular value decomposition (SVD).** Linear models serve as the basis of all continuous gene-network approaches currently available to model typical time-course gene-expression data sets (see [16] for a review). These data sets consist of M pairs of input-output states, represented by D = **{X, Y}.** Matrix **X** is the *N* × *M* gene-expression matrix at time *t*. The columns of matrix **X** labeled by index *v*, x^*v*^, correspond to the experiments, while the rows indicate individual genes. The same is valid for the gene-expression matrix at time *t* + **Δ***t*, **Y**. For a given D, the linear model must map each gene-expression state to the consecutive state, i.e.:

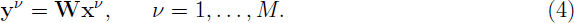

Therefore, in order to find the connectivity matrix, the predicted states from a given input state *x*^*v*^ of the training set must be as close as possible to the output state *y*^*v*^. An alternative would be to minimise the cost function 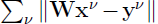. A particular solution with the smallest *L*_2_ norm is given in terms of the SVD of matrix **X**^*T*^ (where superscript *T* denotes the transpose matrix), i.e. **X**^*T*^ = **U·S·V**^*T*^, where **U** is a unitary *M* × *N* matrix of left eigenvectors, **S** is a diagonal *N* × *N* matrix containing the eigenvalues {*s*_1_,…, *s*_*N*_}, and **V** is a unitary *N* × *N* matrix of right eigenvectors [14,17]. Thus, the solution with the smallest *L*_2_ norm represented by **W**_*L*_2__ is given by:

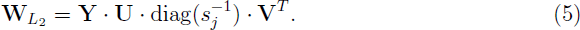

Without loss of generality, all *s*_*j*_ elements whose value is different from 0 were listed at the end of diagonal matrix **S**, and the 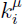 values in Eq. (5) were considered to be 0 if *s*_*j*_ = 0.

The smallest *L*_2_ norm solution cannot be unique. Assuming that x*v* are linearly independent, finding the unique solution requires that M ≥ N. Unfortunately, the inverse problem in GRN involves M ≪ N. Thus, the problem tends to be severely underdetermined, and many solutions can then be consistent with data D. Therefore, all the possible connectivity matrices that are consistent with Eq. (4) can be written in a closed form as:

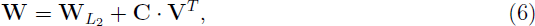

where C is an *N* × *N* matrix whose elements *c*_*ij*_ are 0 as long as *s*_*j*_ **≠** 0. Otherwise, they are arbitrary scalar coefficients. As it will be seen later, the degrees of freedom due to this arbitrariness can be exploited to our benefit [14]. The solution offered by Eq. (5) is implemented to embed the four steady states of *T. cruzi* into the dynamics of the model, without considering the transitions between the states. Eq. (6) is used to uncover the environmental cues by means of using the information provided by the transitions between the different stages of the parasite’s life cycle, and the connectivity matrix associated with the steady states inferred in the previous step.

**Embedding the steady states.** In order to infer the parameter values of Eq. (4) that allow the model to display the same set of steady states as the ones seen in the parasite, we have constructed a training set of size *M*, represented by *D*_*ss*_. Different noise realisations associated with the four stages were added to the steady states. Thus, the columns of matrix **X** are given by:

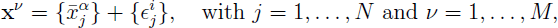

where *α* = 1, 2, 3, 4, and the superscript *i* denotes the noise realisations. In this work, we have used 40 noise realisations for each steady state. Thus, *M* = 4 × 40 = 160. *∊*_*j*_ is a Gaussian noise with mean equal to 0 and a small standard deviation (set at 1% of the expression data). The columns of matrix 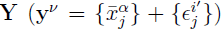 are defined in the same way. We have expanded the size of the training set, thereby making the solutions more robust against the fluctuations. This simple concept is similar to adding a Tikhonov regularisation term in the optimisation process, which has been studied in several neural network problems [18,19]. The constructed training set implies that if at a given time the system is very close to one steady state, it will remain close to that steady state in the next time-step as well (Fig. 2c).

In order to discriminate if an estimated matrix element should be 0 or another reliable value different from 0, we have constructed not only a training set, *D*_*ss*_, but an ensemble of training sets by means of using different noise realisations. For each training set we have computed the minimal *L*_2_-norm solution. Different noise realisations give slightly different solutions. Thus, the ensemble of solutions defines a probability distribution for each weight, *P*_*i,j*_(**ω**). We then performed a location test for each distribution *P*_*i,j*_(*ω*), as illustrated in Fig. 2d. This step consists of testing the hypothesis stating that the true mean value of *P_i,j_* (*ω*) differs from 0 at some magnitude (set at 0.0075). If the p-value associated with this test is greater than 0.01, the hypothesis is rejected, and *w*_*i,j*_ is assigned *p*-value. Otherwise, *w_i,j_* is assigned the mean value of *P*_*i,j*_(*ω*). This procedure allows us to obtain a sparse connectivity matrix, **W_ss_**, that is compatible with the steady states.

**Figure 2:**
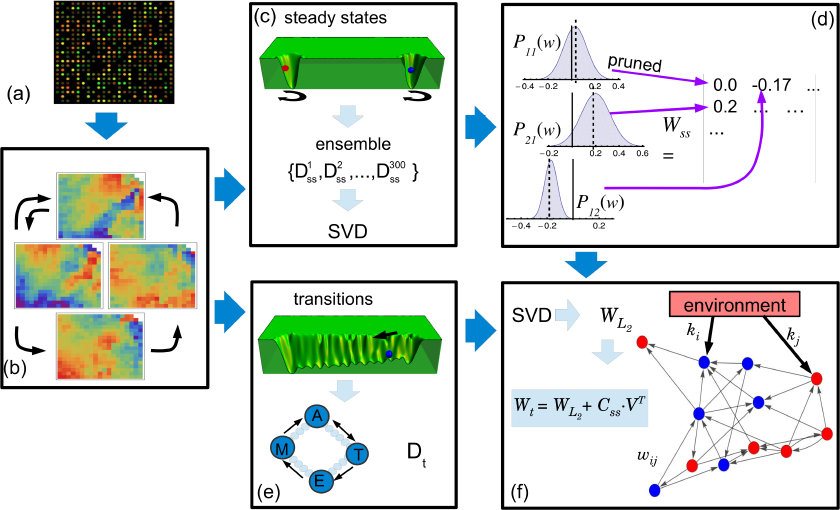
Schema of the network inferring method. (*a*) The microarray data corresponding to the parasite’s four steady states are normalised. (*b*) The total of 8,904 gene-expression levels of each stage is reduced to 339 clusters representing the variables of our systems. (*c*) An ensemble of 300 training sets including fluctuations around the steady states is constructed from the steady states. Using singular value decomposition (SVD), the minimal *L*_2_-norm solution for each **D*_*ss*_* is determined. *(d*) A sparse connectivity matrix, **W**_*ss*_, is derived from the probability distribution Pij (*ω*) by using a pruning method based on a location test. (e) A new training set is constructed from the transitions between the amastigote (A), epimastigote (E), metacyclic tryp. (M) and trypomastigote (T) stages. Intermediate states (small circles) between the stages are assumed to exist. It is also considered that an unknown external cue (black arrow) is responsible for the transitions. (*f*) By means of using SVD, the *L*_2_-norm solution, **W_L_2__**, is determined. This solution is in turn used to find another solution, **W_t_**, which includes information concerning the steady states. This procedure is used to infer the weighted links between genes, *w*_*i,j*_, and to answer two questions: which genes are affected by the external cues, and how they are regulated (up or down) by the environment.

**Embedding the transitions between the steady states.** In order to extend our analysis by including the environmental cues, we have used the extended versions of **W** and *x* described by Eq. (3). To embed the transitions between the steady states, we have considered that these transitions occur gradually and through the shortest possible path between the steady states. Thus, if the system is in the steady state x^*α*^ and is driven to the steady state x^*β*^ due to an external cue *μ* = *β*, then the system performs a series of small transitions between intermediate states represented by *x*^α,β^ (*t*). These intermediate states were constructed by means of a linear combination of the initial and final steady states, i.e. *x*^α,β^ (*t*) = ((*n*_*i*_ − *t*) x^α^ + *t* x^β^)/*n*_*i*_ with t = 0,1, 2,…, *n*_*i*_. As it can be seen, x^*α,β*^(0) and x^*α,β*^(*n*_*i*_) coincide with the steady states x^*α*^ and x^*β*^, respectively. Thus, using these intermediate states, we constructed a new training set *D*_*t*_, where the columns of matrices **X** and **Y** are defined as follows:

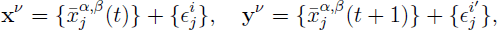

where t = 0,1, 2,…, *n*_*i*_-1. We have used *n*_*i*_ = 10, which implies 10 small transitions. The pairs (**α, β**) correspond to the allowed transitions between the steady states; five transitions in the case of *T. cruzi.* Again, ej is a Gaussian noise with mean equal to 0 and a small standard deviation (set at 1% of the expression data). In all, four different noise realisations were used, and the size of our training set, *D*_*t*_, was in turn *M* = 200. We then computed its smallest **L*_2_* norm solution, **W***_L_2__*. However, since M < N, this solution was not unique. In order to find a particular solution as close as possible to connectivity matrix **W**_*ss*_, we used Eq. (6) and computed the elements of matrix **C**_*ss*_ that obey the following equation:

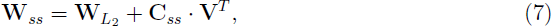

where matrix **W**_*ss*_ was padded with 0 values because **W**_*L*_2__ includes four additional rows and columns corresponding to the environmental cues that are not present in **W**_*ss*_. Eq. (7) is an overdetermined problem that can be solved by applying the interior point method for *L*_1_ regression [14]. The resulting *c*-values were then used to compute a new connectivity matrix (Fig. 2f). This matrix, represented by **W**_*t*_, is not only consistent with the information of the environmental cues and transitions included in *D*_*t*_, but it is also close to **W_ss_.**

## 3 Results

### 3.1 GRN modeling

Key decisions in modeling a gene network system include the choice of variables and the mathematical framework for representing the system dynamics. In this sense, several regulatory network approaches such as Bayesian networks [20], Boolean networks [21], and linear models [13,15, 22] have been suggested. The model must be chosen based on the available data and the ability to infer accurate-enough parameters. The more detailed the model, the more experimental data required to make it work. For instance, when choosing a linear model, in which the expression levels of *N* genes at time *t* determine the changes of such expression levels at time *t* + **Δ***t*, the transition matrix must be computed from *N* pairs of input-output data.

In this work, we have assumed that the system’s state is represented by x(*t*) –the *N*-dimensional vector corresponding to the expression levels of *N* gene clusters measured at time *t*. The GRN dynamics is modeled by a first order Markov model, where the future state depends linearly on the present state and on external perturbations. Mathematically, it is defined by the following equation:

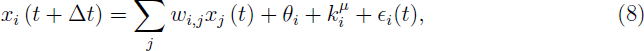

where *w*_*i,j*_ are the elements of the weighted connectivity matrix **W**, and indicate the type and strength of the influence of gene *j* on gene *i* (*w*_*ij*_ > 0 indicates activation, *w*_*ij*_ < 0 indicates repression, and 0 indicates no influence). *θ*_*i*_ is a constant bias term to capture the activity level of gene *i* in the absence of regulatory inputs. We have also added a term indicating the influence of unknown external perturbations, or environmental cues; 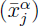, which is the influence of the environmental cue *μ* on gene *i*. Finally, **∊**_*i*_(*t*) is a noise term assumed to be Gaussian with mean 0. The next task in our work was to determine which nodes were affected by external cues –even if those cues were unknown-, and how they were affected. To this end, we considered not only the expression-profile data set information (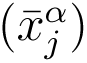), but also some *a priori* information associated with the following biological facts: (i) the parasite’s life cycle has four stages, each of them associated with a measured steady state; (ii) each steady state exhibits some level of noise or fluctuations; and (iii) there are five possible transitions between these four stages. We have assumed that these transitions are the result of different environmental cues acting on certain nodes of the network. Following these facts, we implemented a two-step reverse engineering protocol sketched in Fig. 2. First, we focused on embedding the four steady states into the dynamics of the model, regardless of the transitions between these states. Second, we concentrated on uncovering the environmental-cue effectors considering the transitions between the parasite’s life cycle stages, while using the same connectivity matrix derived in the previous step.

### 3.2 Modeling the steady states of *T. cruzi*

In order to infer connectivity matrix **W**, we have considered the linear model (Eq. (8)) without external perturbations, and have applied the singular value decomposition (SVD) procedure over a training set, *D*_*ss*_, constructed as indicated in Methods. As a result, the dynamical system, together with the derived matrix, has four basins of attraction which correspond to each of the parasite’s stages. This means that whenever the system is in a given basin, it will remain inside that basin as long as there are no external perturbations. Fig. 3a depicts two trajectories (black lines) that illustrate the dynamics of our model in the space spanned by the three principal components. This plot shows that the trajectories fluctuate around the epimastigote and trypomastigote stages. The other two stages, amastigote and metacyclic tryp., showed similar behavior (data not shown). Fig. 3b depicts the time course of the overlap between the state of the system at time t and the epimastigote stage, and the overlap between the state of the system at time t and the trypomastigote stage. Fig. 3c shows a 2D schematic illustration of the pseudo-potential landscape with the four basins of attraction.

The elements of matrix **W** are continuous variables and, consequently, they are associated with not-null values. However, the statistical analysis of known regulatory networks has revealed that such networks have a sparse nature, i.e. the number of actual edges in a network is very small compared to the number of possible edges [23,24]. Such sparsity is difficult to obtain when dealing with continuous weights. Thus, the inferred matrix elements at the end of the reverse engineering process should be either 0 or another reliable value different from 0.

**Figure 3:**
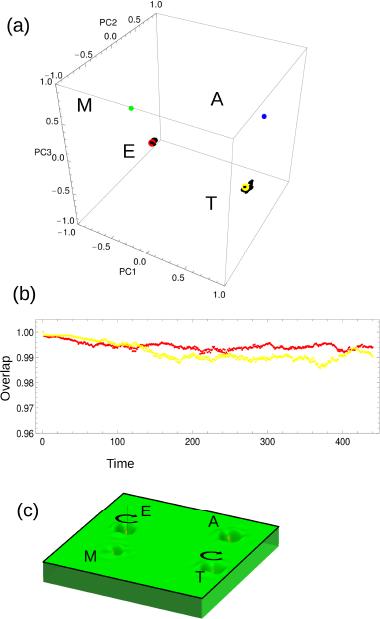
Stability of the steady states. (*a*) The plot shows the positions of the four steady states of the parasite’s life cycle in the space spanned by the three principal components. The black trajectories around the epimastigote and trypomastigote stages are the result of simulations conducted using the model (Eq.(8)) without external cues. A slightly perturbed steady state was used as the initial condition. The system fluctuates around the corresponding steady state. The amastigote and metacyclic tryp. stages showed similar behavior (data not shown). (*b*) Temporal behavior of the overlap between the state of the system at time *t* and the epimastigote steady state (red) or the trypomastigote steady state (yellow). (*c*) 2D projection of the pseudo-potential landscape with the four basins of attraction corresponding to each of the parasite’s stages. The circular black arrows represent the system’s fluctuation around the steady states, just as seen in figure 3a.

In the spirit of inferring a sparse weight matrix that allows the system to display the four steady states, we have used a kind of bootstrap method. To this end, an ensemble of 300 training sets was constructed by means of adding different noise realisations to the steady states, as described in Methods. Using SVD we computed a solution for each training set, obtaining a probability distribution for each weight, *P*_*i,j*_(*ω*). The next step was to assign a value to each element of the connectivity matrix, while carefully assessing the significance of the weight values. We performed a location test to prune the non-significant weights, as illustrated in Fig. 2d, and constructed a sparse connectivity matrix, **W**_*ss*_, which supports the data set. At the significance level of 0.01 there are 11,470 links between genes, i.e. around 10% of the elements of **W**_*ss*_ are not null. Even with this average node degree, the visualisation of the resulting network poses a challenge. In order to overcome this difficulty, we have displayed only a small fraction of the nodes (around 470 links with *p*-value less than or equal to 10^−200^). Fig. 4 shows the GRN. The two weakly connected subnetworks seen in the graph reveal a modular organisation of the network at the significance level used. As it will be seen later, one of these subnetworks is linked to the parasite’s life cycle. The 11,470 links (all not-null elements of the matrix) between genes are listed in S4 Table.

Extracting valuable information from a network made of 10,000 links is a complex task. One way to overcome this problem is by considering only the more important regulators of each steady state. Since the whole regulatory output of a gene depends on the gene’s activity level, some genes can be important regulators in one state, while their activity level in the other three states is low (i.e. *x_i_* ∼ 0). With this in mind, we have constructed network plots that emphasise the most important links in each steady state; that is to say, those links with 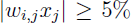 of |*x_i_*|. The plots in S2 Fig. depict the link-derived networks for each of the parasite’s four stages: amastigote (S2a Fig.), epimastigote (S2b Fig.), metacyclic tryp. (S2c Fig.), and trypomastigote (S2d Fig.). As it can be seen, some clusters present regulatory activity only in one particular state. For example, clusters 302, 308 and 333 only appear as relevant regulators in the metacyclic tryp. state, the epimastigote state and the trypomastigote state, respectively. Other clusters, however, are important regulators in all four steady states, as is the case of clusters 326, 336 and 337. Detailed biological information about the genes belonging to the more relevant clusters is listed in S5 Table. After analysing the data obtained for each of *T. cruzi*’s four stages, we have found 47 clusters with important regulatory activity. These clusters include a total of 68 genes: 25 encoding uncharacterised proteins, and 43 coding for proteins with known functions. Among the latter, the most abundant proteins are *trans*-sialidase (TS) (encoded by nine different genes), amastin (encoded by five different genes), and mucin TcMUCII (encoded by four different genes).

**Figure 4:**
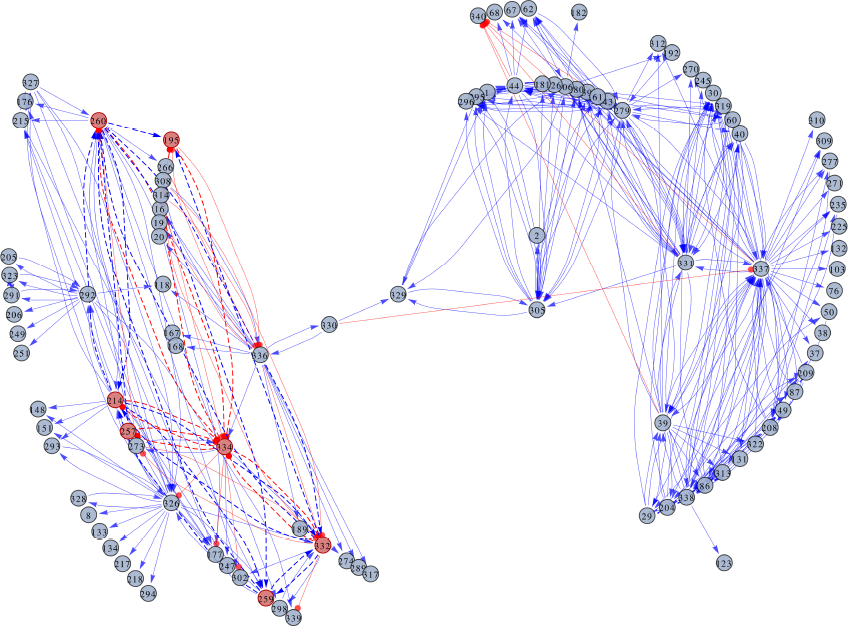
GRN representation of the steady states of *T. cruzi*. The network edges represent the regulatory links between the gene clusters, while the nodes represent the clusters themselves. The labels inside the nodes correspond to the cluster IDs. Additional information about the clusters can be found in tables S5 and S7. The regulatory links indicate either the activation (arrows) or the repression (lines ending in circles) of the clusters. A seven-node subnetwork that controls the dynamics of the parasite’s life cycle is highlighted.

Besides the main four basins of attraction linked to the known steady states displayed in Fig. 3, the system dynamics might include other basins of attraction not-linked to known phenotypes. An exhaustive search for these spurious attractors was performed, and another 20 small attractors where the system can be trapped were found. Fortunately, these new basins of attraction disappeared once subjected to the effects of external cues.

### 3.3 Modeling the phenotypic transitions of *T. cruzi*

After embedding the steady states of the parasite into the GRN dynamics, our analysis was extended to include the transitions that take place between those states as a result of environmental cues. The fact that these transitions in the presence of a given external perturbation occur gradually was taken into account. Since no data about the intermediate states between the steady states are available, we have constructed a training set, represented by *D_t_*, considering that the system performs transitions between the initial and final steady states through the shortest possible path. For details about the construction of the training set, see Methods. As this training set has M < N, there exist infinite solutions compatible with *D*_*t*_. We have chosen the closest solution to the connectivity matrix that uses nothing but the steady states information, i.e. the closest to **W**_*ss*_. Thus, our connectivity matrix is represented by:

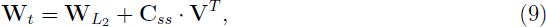

where **W**_*L*_2__ is the corresponding minimal **L*_2_*-norm solution obtained by SVD for *D*_*t*_. Matrix *C*_*ss*_ was computed by the interior point method as described in Methods. The new connectivity matrix, **W**_t_, is consistent with the information of the environmental cues and transitions included in training set *D*_*t*_. As **W**_*t*_ is also very close to **W**_*ss*_, it consequently inherits the ability to support the multi-stability of the parasite’s life cycle.

In order to test the ability of the model (Eq. (8)) to emulate the observed dynamical behavior, simulations under different external cues were performed. Each of these simulations was performed considering that the system is initially in one of the parasite’s steady states, and that an external cue *μ* is acting. The simulations were performed by running 12 iterations of the model (Eq. (8)), and recording the system’s state at each of these 12 steps. The temporal evolution of the 339 variables of the system was compiled in movies available as S1-S5 Movies. S1 Movie shows the simulated phenotypic transition from the amastigote stage to the trypomastigote stage when external cue *μ* = 4 is acting. S2-S5 Movies, on their part, illustrate the modeling results of the remaining phenotypic transitions. In all cases, the final state of the system is in agreement with the expected state regarding the acting external cue. This agreement can be better appreciated when using a principal component analysis procedure for dimension reduction. Fig. 5 depicts a set of trajectories corresponding to four of the five phenotypic transitions in the 3D space spanned by the main principal components. There are 20 alternative trajectories for each simulated phenotypic transition. All of the trajectories for a given transition have the same initial condition, are affected by the same external cue, but present particular noise realisations. Hence, it can be said that the model is able to reproduce the dynamics of *T. cruzi*’s life cycle.

**Figure 5:**
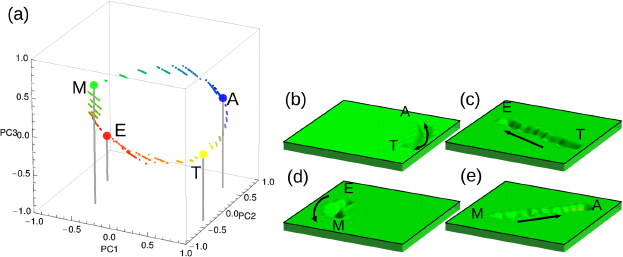
Representation of transitions between the steady states caused by external cues. (*a*) The plot shows the trajectories of the system from an initial to a final steady state under the influence of an external cue in the space spanned by the three principal components. A slightly perturbed steady state was used as the initial condition. Since amastigote-to-trypomastigote and trypomastigote-to-amastigote transitions overlap, only the first one is shown. Each trajectory has 10 intermediate states represented by small circles. (*b*), (*c*), (*d*) and (*e*) 2D projections of the pseudo-potential landscapes corresponding to the phenotypic transitions mentioned above.

In our model, the phenotypic transitions are caused by an environmental cue *μ* through parameter 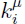, i.e. the gene clusters associated with large positive (or negative) 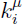 values are activated (or inhibited) by the acting external cue *μ*. k^*μ*^ values are distributed around 0. In order to identify the key connections that modulate the network behavior under external cues, we have selected those gene clusters with k^*μ*^ values greater (lower) than the 95th (5th) percentile of the distribution. These clusters are listed in S6 Table. A total of 166 externally regulated genes belonging to 86 different clusters were found. While 73 of these genes encode uncharacterised proteins, the other 93 genes code for proteins with known functions. Just as in the steady states, the most abundant proteins are TS (encoded by 21 different genes), amastin (encoded by six different genes), and mucin TcMUCII (encoded by five different genes). The difference, however, is that when considering the transitions, these proteins act no longer as regulators, but they are up-or down-regulated instead. According to their GO annotations, all of the TSs have exo-alpha-sialidase activity. Uncharacterised proteins without GO annotations have been analysed using the InterproScan software [25], and the results are summarised in S7 Table. According to this analysis, 39% of these proteins are membrane proteins. We have also analysed the data corresponding to each type of host separately. In this sense, we have found that those genes affected by the external cues leading to the parasite’s two mammalian stages are inversely regulated, while those genes affected by the external cues leading to the parasite’s two insect stages are equally regulated.

The next step in our analysis was to identify the module that controls the parasite’s life cycle. This is a difficult task because it involves the isolation of a small subset of nodes and regulatory connections out of a network of 10,000 links. In principle, the number of possible subnetworks within a network of such a size is very large. Consequently, the evaluation of the subnetworks’ dynamics is not possible. In order to solve this problem, we reduced the search space. With this in mind, we considered only those circuits that involve nodes with important regulatory roles. To this end, we have used the list of regulatory clusters shown in S5 Table, and have written a script to search for cyclic graphs, i.e. closed loops, containing such nodes in matrix **W**_*t*_. With this set of modules, we then searched for those subnetworks with the ability to emulate the parasite’s dynamics. At this point, our model had to be simplified. In order to evaluate the dynamics of the system, we have considered that variable xi is a Boolean variable, and that the system’s evolution is given by:

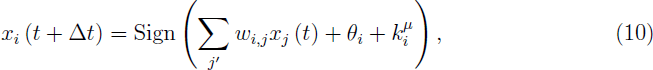

where index *j*’ in the sum runs only over the nodes belonging to the module under evaluation. The parameter values are taken from **W_t_**, and listed in S8 Table.

As a result of the searching process, we were able to identify a seven-node module, containing a total of nine genes. Fig. 6a illustrates the architecture of this subnetwork.

The seven clusters forming the subnetwork are: 195, 214, 257, 259, 260, 332, and 334. Relevant information about these clusters and their composing genes is shown in Table 1. Three of the nine genes code for uncharacterised proteins (Q4DTV8, Q4DVU8 and Q4E589). According to their GO annotations, Q4DTV8 has hydrolase activity (acting on carbon-nitrogen but not peptide-bonds, in linear amidines), Q4DVU8 has transporter activity, and Q4E589 has catalytic activity. The other six genes code for: an hexokinase (Q4D3P5), a **δ**-1-pyrroline-5-carboxylate dehydrogenase (Q4DRT8), a quinone oxidoreductase (Q4DHH8), a glutamate dehydrogenase (Q4DWV8), a peptidyl-prolyl cis-trans isomerase (Q4E4L9), and a metaciclina II (Q4E2M3).

**Figure 6:**
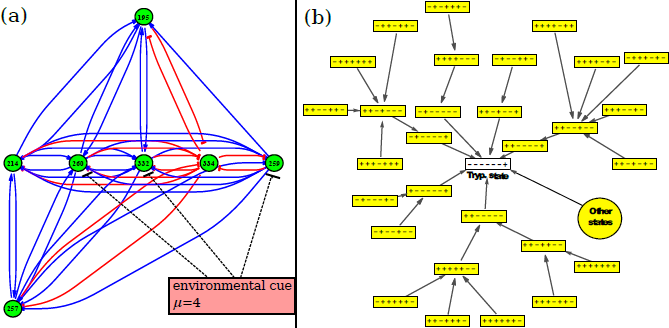
Life cycle module. (*a*) Architecture of the seven-node subnetwork that controls the dynamics of the parasite’s life cycle. The action of environmental cue *μ* = 4 is shown as an example. (*b*) Boolean dynamics of the life cycle module. The basin of attraction of the seven-node module under the action of environmental cue *μ* = 4 is shown. This external signal leads the network to the trypomastigote state. Here, the nodes represent the module states and the edges represent the transitions. The module states are characterised by the sign of the clusters, which in turn are arranged in box according to their cluster IDs. Under the action of this perturbation, the final state is always the trypomastigote stage (white box). Some states reach this final state by going through different intermediate steps, while others (represented by the biggest circle) reach it in only one step.

The identified subnetwork reproduces many important dynamical features observed in the life cycle of *T. cruzi.* On the one hand, the phenotypic transitions from epimastigote to metacyclic tryp., from amastigote to trypomastigote, and from trypomastigote to epimastigote are reproduced under the influence of the corresponding external cue. And on the other, the phenotypic stages epimastigote, metacyclic tryp., and trypomastigote correspond to steady states of the subnetwork’s dynamics. As an example of the subnetwork’s dynamics, Fig. 6b illustrates the basin of attraction of the module under the action of environmental cue *μ* = 4. This external signal leads the network to the trypomastigote state. The figure shows that regardless of the initial state (there are 128 Boolean states), the final stop of the trajectories in the Boolean space is always the trypomastigote stage. Similarly, when the environmental cue is *μ* = 2 or *μ* = 3, the obtained basin of attraction is the epimastigote or the metacyclic trypomastigote stage, respectively.

**Table 1.** Subnetwork information.

| Cluster ID | Gene ID | Uniprot ID | Putative function | $\mu$ | $\mathbf{k}^\mu$ coefficient |
| --- | --- | --- | --- | --- | --- |
| 195 | 6382 | Q4DTV8 | hydrolase activity | 3 | -0.1805 |
|  | 6588 | Q4D3P5 | hexokinase | 3 | -0.1805 |
| 214 | 121 | Q4DRT8 | delta-1-pyrroline-5-carboxylate dehydrogenase | 2 | 0.4251 |
| 257 | 519 | Q4DHH8 | quinone oxidoreductase | 2 | 0.2256 |
| 259 | 8762 | Q4DVU8 | transporter activity | 2 | 0.3033 |
| 260 | 4053 | Q4E589 | catalytic activity | 2 | 0.3479 |
|  | 5564 | Q4DWV8 | glutamate dehydrogenase | 2 | 0.3479 |
| 332 | 4595 | Q4E4L9 | peptidyl-prolyl *cis-trans* isomerase | 3 | -0.304 |
| 334 | 1518 | Q4E2M3 | metaciclina II | 2 | -0.2685 |

List of the clusters and genes forming the parasite’s life cycle subnetwork, the protein function, the most relevant external cue, and the corresponding values of $\mathbf{k}^\mu$ coefficients needed to reproduce the dynamical features of the system.

## 4 Discussion and Conclusions

One fundamental open question in systems biology is how cells that share the same genome exhibit notably different gene-expression patterns or distinct phenotypes. This question is closely related to the process of establishing cell fates during development. A widely used picture to describe these phenomena is Waddington’s epigenetic landscape, a phenomenological metaphor which corresponds to an energy landscape with many local minimums where the system moves regardless of whether environmental cues are present [26-28].

Despite the simplicity and elegance of Waddington’s concept, it lacks quantitative mechanistic details. Given the significance of a quantitative understanding of cell phenotypic transitions, many efforts have been made to develop predictive mathematical frameworks [28-32]. Although some advances have been made for low dimensional systems, the application of these mathematical frameworks to higher dimensional models remains a theoretical challenge.

In this work, we have developed a reverse engineering approach to identify the gene network structures responsible for the observed dynamical properties of a high dimensional biological system. These dynamical properties include the steady states associated with the stable phenotypes, and the phenotypic transitions observed in *T. cruzi*’s life cycle. We have assumed that each of the five phenotypic transitions occurs in response to the external cue corresponding to the final state of the transition. For this reason, we have modeled the life cycle of *T. cruzi* as if it were an open dynamical system. Our methodology for embedding the observed expression patterns into the GRN dynamics adds several new ingredients such as the use of an ensemble of noise-perturbed training sets, and a pruning procedure to identify the significant network links. The information of the transitions between the stable phenotypes was used to develop an optimisation procedure. This reverse engineering procedure has been successfully used to identify one key network module that explains three of the five phenotypic transitions.

Besides the development of a model with the ability to emulate the parasite’s dynamics, the information presented in this work (S5 and S6 Tables) could be useful to assign previously unknown putative functions to some genes. In this sense, our results suggest that amastin genes could act as key regulators. This finding is consistent with a previous study in which it is shown that amastin may increase *T. cruzi*’s differentiation rates both in the insect and in the mammalian hosts [33]. On the other hand, our finding of TS-coding genes acting as regulators in the amastigote stage adds relevant information to the resulting parasite state. Furthermore, we have found that these same TS-coding genes are inhibited in the transitions leading to the amastigote stage, and activated in the transitions leading to the trypomastigote stage. Considering that TS plays a key role in *T. cruzi*’s infectivity and that this enzyme is not present in mammals, TS constitutes a potential target for the development of novel drugs to treat or prevent Chagas disease [34-36]. Finally, we have found four mucin genes belonging to the TcMUC family that act as regulators both in the mammalian and in the insect parasite’s stages. It is known that this family of mucins is expressed only in the mammalian stages [37].

The present approach could be adapted to and useful for better understanding other single cell parasites with multiple developmental stages such as *T. brucei, P. falciparum* and *Leishmania.* Uncovering the core circuit that underlies the dynamics of these parasites’ life cycles could open the door to new possibilities: the development of applications to reprogramme the parasites’ life cycles, and the finding of new therapeutic targets against the parasites.

## Authors’ contributions

AC participated in data analysis and interpretation, participated in the design of the study and drafted the manuscript; LD conceived of the study, designed the study, participated in data analysis and interpretation, and drafted the manuscript. All authors gave final approval for publication.

## Acknowledgments

The authors wish to thank Geoffrey Siwo for his critical review of the manuscript. A.C. is a postdoctoral fellow of the CONICET (Argentina). L.D. is a research member of the CONICET (Argentina).

## Supporting Information

**S1 Fig. Clustering and dimension reduction.** Davies-Bouldin index (DBI) as a function of the number of clusters, *N_c_*, used in the clustering procedure. The arrow in *N_c_* = 339 indicates the optimal number of clusters used in subsequent procedures.

**S2 Fig. Main regulatory clusters for each steady state.** The plots represent the networks derived from amastigote (a), epimastigote (b), metacyclic tryp. (c), and trypomastigote (d) stages.

**S1-S5 Movies. Transitions between the parasite’s stages.** The animated matrix plots show the evolution of the system from one of the initial phenotypic stages shown in Fig 1b. The movies correspond to different system evolutions: from the amastigote stage under the action of *μ* = 4 (S1 Movie), from the trypomastigote stage under the action of *μ* = 2 (S2 Movie), from the epimastigote stage under the action of *μ* = 3 (S3 Movie), from the metacyclic tryp. stage under the action of *μ* = 1 (S4 Movie), and from the trypomastigote stage under the action of *μ* = 1 (S5 Movie). Each movie is composed of 12 frames; one for each step in the simulation. All simulations show a clear similarity between the states associated with the last two frames and the corresponding target stage indicated by *μ*.

**S1 Table. Gene-expression profile of *T. cruzi*’s life cycle.** Log-norm expression levels corresponding to 8,904 *T. cruzi* genes, obtained from microarray experiments as indicated in Methods. The gene IDs listed in the first column correspond to our own gene numbering. The second column lists the microarray oligo IDs. The last four columns correspond to the gene-expression levels in each stage of the parasite’s life cycle: amastigote, epimastigote, metacyclic tryp. and trypomastigote, respectively.

**S2 Table. Cluster composition.** Each row corresponds to one cluster ID and lists all genes (gene IDs) belonging to that cluster.

**S3 Table. Intra-cluster averages of the expression levels.** Gene activity levels of each cluster (rows) in each different stage (columns) used in all further modeling computations.

**S4 Table. Regulatory links.** List of the 11,470 significant weights (at a significance level of 0.01) needed for the maintenance of the parasite’s steady states. The first column indicates the regulatory cluster IDs. The second column indicates the regulated cluster IDs. The third column lists the mean values of each cluster, averaged over the ensemble of 300 training sets. The fourth column lists the associated standard deviations. The last column lists the p-values (the probabilities of the location test).

**S5 Table. Main regulatory genes**. List of the genes with important regulatory activity for the maintenance of the parasite’s steady states. The steady states are indicated by a capital letter: amastigote (A), epimastigote (E), metacyclic tryp. (M) and trypomastigote (T).

**S6 Table. Main regulated genes.** List of the genes regulated by the external cues responsible for the transitions between the steady states.

**S7 Table. Analysis of uncharacterised proteins.** Result of the Interproscan analysis of the uncharacterised proteins without GO annotations, listed in S5 and S6 Tables. Uncharacterised proteins with no Interproscan information are listed at the end.

**S8 Table. Regulatory links of *T. cruzi*’s life cycle subnetwork.** Estimated values of parameters *ω_i,j_*, **θ**_*i*_, and *k*^μ^_*i*_, extracted from matrix **W**_*t*_.

**Abbreviations:** GRN, gene regulatory network; SVD, singular value decomposition; TS, *trans*-sialidase

